# Critical roles of ERK and Akt signaling in metabolically dependent memory formation in *Drosophila* larvae

**DOI:** 10.1101/2024.06.28.601201

**Authors:** Hanna R. Franz, Annekathrin Widmann

## Abstract

In the fruit fly *Drosophila*, memory formation is intricately linked to metabolic state. Our previous findings showed that the preferential support of memory under increased energy levels is mediated by the insulin receptor in the mushroom body (MB), a key memory center in the insect brain. This study analyzed the role of the insulin receptor’s downstream pathways, Ras-Raf-MEK-ERK and PI3K/Akt, in metabolically dependent memory formation. We evaluated their impact on memory processes by employing RNA interference-mediated downregulation of key components, extracellular signal-regulated kinase (ERK) and protein kinase B (Akt), in the MB of *Drosophila* larvae. To enhance energy levels, larvae were fed sugar for 60 minutes prior to aversive olfactory conditioning. Our findings revealed that genetically downregulating ERK and Akt in the MB significantly impaired metabolically dependent memory formation, without affecting naive sensory perception or olfactory learning. Immunohistochemical analysis confirmed the presence of active forms of ERK and Akt in the MB, underscoring their roles in modulating memory processes under elevated energy levels. The pivotal roles of these kinases suggest a broader implication of insulin signaling in memory formation under different metabolic conditions, and illuminate the connections between metabolic regulation and memory dynamics in the MB.

## Introductions

Learning and memory, which are critical for the survival and adaptation of most organisms, encompass both transient, short-lasting memories and robust, long-lasting memories [1–3]. In the fruit fly *Drosophila melanogaster*, aversive olfactory memory can also be categorized into these forms, each of which is established by distinct training protocols [4–7], conserved signaling pathways [8,9], and neuronal activity within the memory circuits [4–10]. These memory processes are further influenced by various factors, including physiological states and behaviors such as sleep patterns, aging, mating [11–20], forgetting [21–25], additional neuronal activities outside the primary learning circuits [26,27], additional signaling pathways [5,28,29], and cellular mechanisms such as proteasomal and lysosomal degradation, glial function, and epigenetic modifications [30–37]. This complexity underscores the need for favorable conditions for robust memory formation, with the energetic state of the organism being a critical factor [38]. For example, in *Drosophila*, aversive olfactory long-term memory (LTM) typically requires *de novo* protein synthesis and spaced training sessions, which involve repeated learning episodes separated by rest intervals [4–6,39]. However, starvation can suppress LTM formation to ensure survival, because it is energetically costly [40,41]. Conversely, when the animal’s metabolic state is elevated, LTM can be formed without the traditional requirement for spaced training sessions [42]. A key mediator of an organism’s metabolic state is insulin signaling, which reflects the nutritional status of an organism. Insulin signaling induces a systemic response that regulates anabolic and catabolic pathways and influences learning and memory in various species [43–47]. More specifically, within the mushroom body (MB), a key site for associative learning and memory in insects [48,49], the insulin receptor significantly influences memory formation in *Drosophila* [15,42,50,51]. Our previous research has shown that the insulin receptor is critical for memory formation under conditions of elevated energy levels, thereby integrating the metabolic state of the organism with memory processes [42].

Our current study extends this line of inquiry to the downstream signaling pathways of the insulin receptor, particularly the Ras-Raf-MEK-ERK and PI3K/Akt pathways [52]. Our goal was to explore the roles of two key components, the extracellular signal-regulated kinase (ERK) and protein kinase B (Akt), in the regulation of memory formation in larvae in an elevated energetic state, reflecting their established significance in memory processes across various species [53]. To achieve this goal, we first performed immunohistochemical analyses, which confirmed the presence of pERK and pAkt in the MB and suggested their potential role in the modulation of memory in response to an increased energy supply. We then utilized RNA interference (RNAi) to specifically downregulate the ERK and Akt in the MB in larvae and examine how this downregulation influences memory formation under elevated energetic conditions. Our results showed that reducing ERK and Akt levels in the MB significantly altered memory formation under conditions of elevated energy states. This effect on metabolically dependent memory formation was evident because naive sensory perception and olfactory learning remained unaffected. This study highlights the critical role of ERK and Akt kinases in metabolically dependent memory formation in *Drosophila* larvae and suggests an intriguing mechanism by which these kinases integrate metabolic regulation with memory dynamics.

## Material and methods

### Fly stocks

Fly strains were reared on standard *Drosophila* medium at 25°C with 60% humidity under a 12-h light-dark cycle. For the knockdown of ERK and Akt, two RNAi effector lines from the Vienna Drosophila Resource Center (VDRC)—*UAS-Akt-RNAi#1* (II) (103703) and *UAS-ERK-RNAi#1* (II) (109108)—and two lines from the Bloomington Drosophila Stock Center (BDSC)—*UAS-Akt-RNAi#2* (III) (33615) and *UAS-ERK-RNAi*#2 (III) (34855) were utilized [54,55]. Driver lines in this study were *elav*-Gal4 (I) and *H24*-Gal4 (III) [56–58]. *H24*-Gal4 and *UAS-ERK-RNAi#2* were outcrossed six times to wild-type *Canton-S*. Genetic controls were established by crossing Gal4-driver lines and UAS-effector lines to wild-type *Canton-S*. All stocks used in this study are listed in Table 1.

### Naïve odor preference and high-salt avoidance experiments

To study naïve olfactory preferences and high-salt avoidance, 30 larvae (6 days old) were placed along the midline of a Petri dish (92-mm diameter Sarstedt, Nümbrecht, 82.1472), marked by a 1-cm neutral zone. For olfactory testing, the Petri dish was filled with 2.5% (w/v) agarose (Sigma Aldrich, A5093). On opposite sides of the dish, two custom-made Teflon containers (4.5 mm diameter) with perforated lids were placed: one contained 10 μl of a specific odor and the other was empty. For the odors, either amyl acetate (AM, Sigma Aldrich, 109584) or benzaldehyde (BA, Sigma Aldrich, 418099) was used; when necessary, these substances were diluted in paraffin oil (Sigma Aldrich, 18512). After 5 minutes, the numbers of larvae present on the odor side (OD), the empty container side (EC), and the neutral zone (NZ), were counted. A preference index (PREF) was calculated by subtracting the number on the empty container side from the odor side, and then dividing by the total number of larvae, as follows:

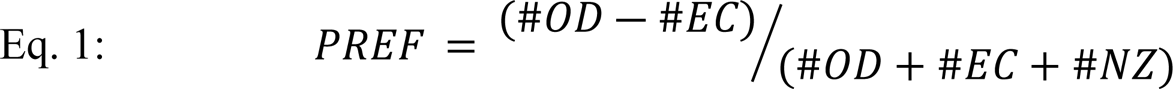

For high-salt avoidance testing, the same Petri dish setup was used, but for this assay one side contained 2.5% agarose and the opposite side contained 2.5% agarose mixed with 1.5 M sodium chloride (salt, Sigma Aldrich, S7653). The neutral zone was filled with 2.5% agarose mixed with 1.5 M sodium chloride. After 5 minutes, the numbers of larvae present on the salt side (SALT), the agarose side (AG), and in the neutral zone (NZ) were counted. An avoidance index (AVOID) was calculated by subtracting the number of larvae on the agarose side from the number on the salt side, and then dividing by the total number of larvae, as follows:

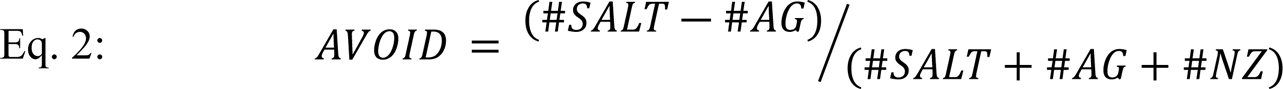

### Aversive olfactory learning and memory

Standard aversive olfactory conditioning experiments were performed using an odor–high-salt conditioning paradigm, as previously described [5]. Briefly, experiments were conducted on assay plates (92-mm diameter) filled with a thin layer of 2.5% agarose containing either pure agarose or agarose plus 1.5 M sodium chloride. Amyl acetate, diluted 1:250 in paraffin oil, and undiluted BA were used as olfactory stimuli, each dispensed in a volume of 10 μl into custom-made Teflon containers (4.5-mm diameter) equipped with perforated lids. For odor–high-salt learning, groups of 30 larvae (6 days old) were exposed to each odor for 5 minutes; one odor was paired with sodium chloride (CS+) and the other was presented without (CS-). Learning and memory were assessed on fresh agarose plates containing 1.5 M sodium chloride, with both odors presented on opposite sides of the plate. After 5 minutes, the larvae were counted. A preference index (PREF) was calculated by subtracting the number of larvae on the CS-side from that on the CS+ side, and then dividing by the total number of larvae. To specifically measure associative learning effects, a performance index (PI) was calculated by averaging the preference indices from two reciprocal experiments, as follows:

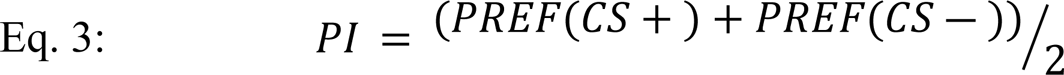

### Sugar feeding

The energetic state of larvae was manipulated as described previously [42]. Briefly, a group of 30 larvae was gently washed out of the standard *Drosophila* food and was fed for 1 hour with 0.15 M sucrose (Sigma Aldrich, 84097) mixed with tap water in a Petri dish (35-mm diameter, Sarstedt, Nümbrecht, 82.1135.500) containing a thin layer of 2.5% agarose at 23°C. Zeitgeber time and humidity were kept constant for these experiments. After being fed, the larvae were washed gently with tap water and transferred to a water droplet.

### Protein extraction

Protein samples were prepared from 20 adult female heads per sample, homogenized in 50 µl of ice-cold lysis buffer (1 mM EDTA, 1:1000 protease-inhibitor cocktail, 1:100 phosphatase-inhibitor cocktail 2, in 1X PBS) using a Qiagen TissueLyser LT for 2 minutes at 50 Hz. After homogenization, samples were centrifuged at 1000 rpm for 15 seconds, and 40 µl of supernatant was collected into chilled Eppendorf tubes. Protein concentrations were determined using the Bradford Protein Assay Kit (Thermo Fisher Scientific, 23200) according to the manufacturer’s protocol, with Coomassie Blue-protein interaction assessed by absorbance at 595 nm using a microplate spectrophotometer (BioTek, Vermont, USA, VT 05404-0998). Sample concentrations were then adjusted to 1000 or 2000 µg/ml for Western blot analysis based on total protein yield. A detailed list of reagents used is provided in Table 2.

### Western blot and quantification

Protein samples were mixed 1:1 with 2X Lämmli sample buffer (Sigma-Aldrich, S3401-1VL) and denatured at 95°C with shaking at 400 rpm for 10 minutes. Subsequently, the samples underwent a series of two-fold dilutions across three steps and were applied to an 8-16% gradient SDS-PAGE precast acrylamide gel (NuSep, NN12-816). Electrophoresis was conducted at 300 V and 40 W for 90 minutes (PEQLAB, 45-1010-i), setting the current at 32 mA for a single gel and 65 mA for two gels. Proteins were then transferred to a nitrocellulose membrane (Amersham, 10600007) using a wet chamber system (PEQLAB, PerfectBlue™ Tank-Electro Blotter) at 70 V, 40 W, and 600 mA for 1 hour. Successful transfer was verified by Ponceau S staining (1% (v/v) acetic acid, 0.1% (w/v) Ponceau S in ddH2O). The membranes were then blocked in 5% (w/v) skim milk powder (Sigma-Aldrich, 70166) in 0.3% (v/v) TBS-Tween 20 (TBST; Sigma-Aldrich, P1379) with gentle agitation overnight at 4°C. Incubation with primary antibodies was performed for two nights at 4°C in 0.3% TBST containing a 1:10 dilution of 5% skim milk powder in 0.3% TBST. This was followed by three washes in 0.3% TBST and incubation with secondary antibodies for one night at 4°C in 0.3% TBST containing a 1:10 dilution of 5% skim milk powder in 0.3% TBST. Protein detection was performed by first washing the membrane three times with 0.3% TBST, then rinsing with TBS, followed by application of Pierce™ ECL Western Blotting Substrate (Thermo Scientific, 32209). Chemiluminescence signals were detected for 20 minutes using a FluorChem FC2 system (Cell Biosciences, Santa Clara, USA). To perform a semi-quantitative analysis of pERK protein levels, band densities were quantified using the gel analyzer tool [59] of ImageJ [60]. Initially, Western blot images were filtered using a 2-pixel median filter to remove background noise. Afterwards, profile plots were generated to quantify peak areas representing the density of respective protein bands. Based on the three, two-fold dilution steps per sample, slopes were calculated and normalized to wild-type protein levels and the internal α-Tubulin housekeeping control as follows:

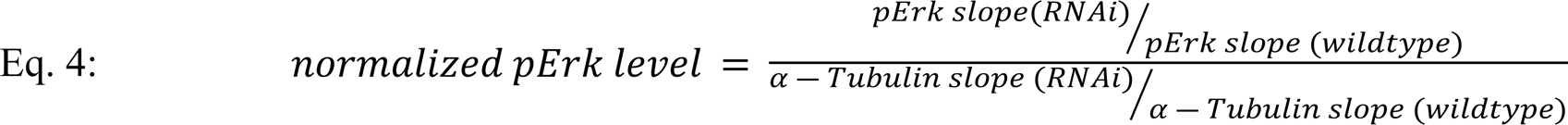

All samples for each experiment were processed on a single Western blot to ensure uniform conditions, and the entire study comprised three independent Western blots. A detailed list of antibodies and reagents used is provided in Table 2.

### Immunohistochemistry of larval brains

Brains from 4-to 6-day-old, third-instar larvae were dissected in ice-cold larval Ringer’s solution with a 1:100 dilution of phosphatase inhibitor cocktail 2 (Sigma-Aldrich, P5726). Tissues were fixed in 4% paraformaldehyde with 1:100 phosphatase inhibitor cocktail 2 in PBS for 60 minutes at room temperature. This was followed by a series of washes in PBST (PBS with 3% Triton^™^ X-100, Sigma-Aldrich, X-100) (once for 10 minutes, twice for 30 minutes), and blocking with 2% (v/v) normal goat serum (NGS) (Thermo Fisher Scientific, PCN5000) in PBST for 90 minutes at room temperature. The brains were incubated in primary antibodies at a 1:25 dilution from a 2% NGS stock solution in PBST, over two nights at 4°C. Subsequently, the samples were washed in PBST at room temperature—twice for 10 minutes and three times for 30 minutes—before being incubated in secondary antibodies at a 1:25 dilution from a 2% NGS stock solution in PBST, for another night at 4°C. A final series of washes in PBST was performed—once for five minutes, once for ten minutes, and twice for 30 minutes. After a brief rinse in PBS, the brains were mounted on a microscope slide within a 6 µL drop of VECTASHIELD Mounting Medium (Vector Laboratories, H-1000) set within a reinforcement ring. Imaging was performed within 48 hours of mounting using a Leica TC SP8 confocal laser scanning microscope. This microscope was equipped with either a Leica Apochromat 20× air objective (NA = 0.70) or a 63× glycerol objective (NA = 1.30). DAPI, Alexa 488, Alexa 568, and Alexa 633 fluorophores were excited at wavelengths of 405 nm, 488 nm, 561 nm, and 633 nm, respectively. The MB was scanned in 1-μm sections along the z-axis, with a zoom factor of 3× and a resolution of 5.28 pixels/μm for 20× scans, and a zoom factor of 1× with a resolution of 5.5440 pixels/μm for 63× scans. Overviews of the central nervous system (CNS) utilized a z-step size of 1 μm, a zoom factor of 1x using ×20 objectives, and a resolution of 1.7600 pixels/μm. All imaging parameters, including laser excitation power, fluorophore detection ranges, zoom factor, and z-step size, were kept consistent across experiments involving different genotypes or treatments. Images were created and analyzed using Fiji [61]. A detailed list of antibodies and reagents used is provided in Table 2.

### Quantification and statistical analysis

All statistical analyses and visualizations were performed using GraphPad Prism 10. Figure alignments and schematic drawings were performed using Adobe Illustrator CC 2022. To compare individual groups against the chance level, we employed Holm-Sidak-corrected, two-tailed, one-sample t tests for normally distributed data (as determined by the Shapiro-Wilk test), and Holm-Sidak-corrected, two-tailed, Wilcoxon signed-rank tests for non-normally distributed data. To statistically analyze differences among groups that met the assumptions of normality (as determined by the Shapiro-Wilk test) and homogeneity of variance (as determined by Bartlett’s test), we conducted a one-way ANOVA followed by Tukey honestly significant difference (HSD) post-hoc pairwise comparisons. In cases where the data significantly deviated from the assumptions of normality and homogeneity of variance, a non-parametric Kruskal-Wallis test was performed, followed by Dunn’s multiple pairwise comparisons. The data for the *H24*-Gal4 control group in the experiments shown in Figs. 3B and 3C, 5B and 5E, 5C and 5F, 5D and 5G, and 6B and 6C were pooled from overlapping experiments. Detailed information on the specific statistical tests used, sample sizes, and descriptive statistics can be found in Tables 3 and 4. For all statistical tests, the significance level was set at *α* = 0.05.

**Fig 1.**
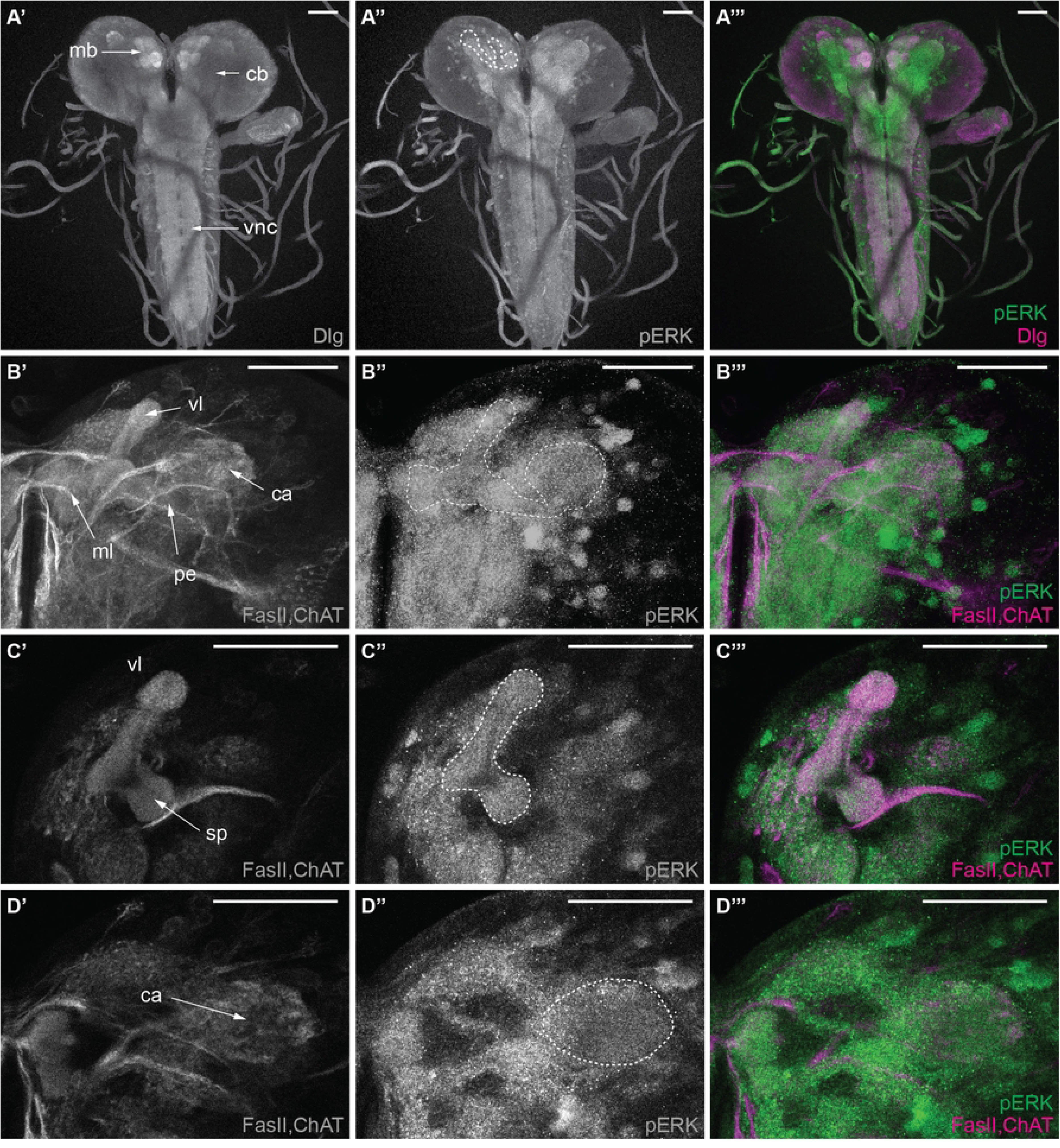
Distribution of phosphorylated ERK (pERK) in the larval central nervous system (CNS) (A’-A’’’) Maximum z-projections of the CNS labeled with anti-Dlg and the phosphorylated antibody anti-pERK. (A’) shows the postsynaptic marker Dlg, (A’’) shows the anti-pERK labeling, and (A’’’) is an overlay of both with Dlg in magenta and pERK in green. (B’-B’’’) Maximum z-projection of the mushroom body (MB) with anti-FasII and anti-ChAT highlighting the neuropil and anti-pERK. (B’) shows the neuropil, (B’’) shows the anti-pERK labeling, and (B’’’) is an overlay of both with neuropil in magenta and pERK in green. The MB calyx, peduncle, and lobes (vl, ml) are outlined in white. (C’-D’’’) Single-slice views highlight the distribution of pERK in MB subregions, including visible portions of the lobes (vl, sp) (C’-C’’’) and calyx (ca) (D’-D’’’), with FasII and ChAT (magenta) labeling the neuropil. Scale bar: 50 µm. Abbreviations: ca (calyx), cb (central brain), ChAT (choline acetyltransferase), Dlg (Disc large), FasII (fasciclin II), mb (mushroom body), ml (medial lobe), pe (peduncle), sp (spur), vl (vertical lobe).

**Fig 2.**
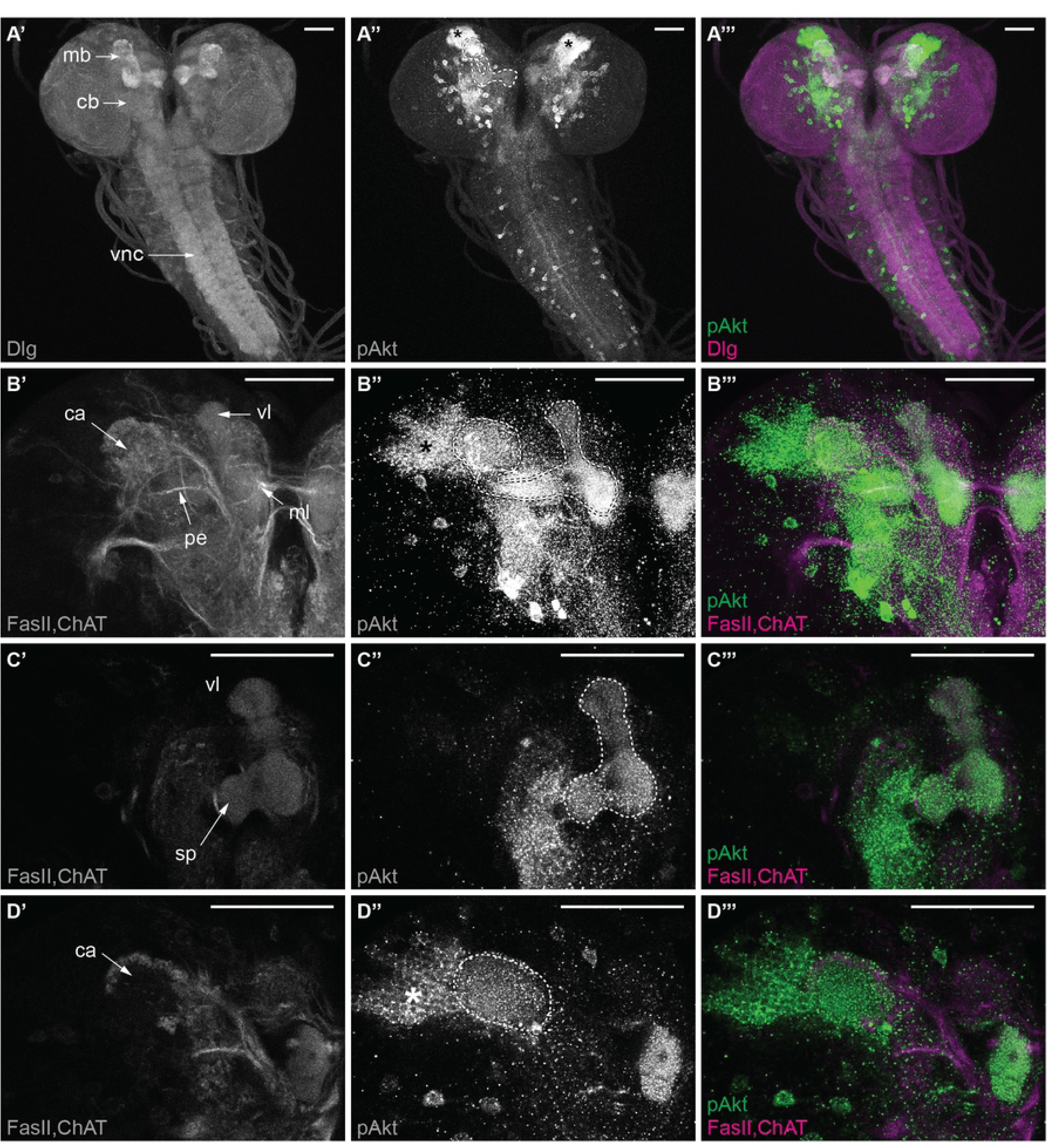
Distribution of phosphorylated Akt (pAkt) in the larval central nervous system (CNS) (A’-A’’’) Maximum z-projections of the CNS labeled with anti-Dlg and the phosphorylated antibody anti-pAkt. (A’) shows the postsynaptic marker Dlg, (A’’) shows the anti-pAkt labeling, and (A’’’) is an overlay of both with Dlg in magenta and pAkt in green. (B’-B’’’) Maximum z-projection of the mushroom body (MB) with anti-FasII and anti-ChAT highlighting the neuropil and anti-pAkt. (B’) shows the neuropil, (B’’) shows the anti-pAkt labeling, and (B’’’) is an overlay of both with neuropil in magenta and pAkt in green. The MB calyx, peduncle, and lobes (vl, ml) are outlined in white. (C’-D’’’) Single-slice views highlight the distribution of pAkt in MB subregions, including visible portions of the lobes (vl, sp) (C’-C’’’) and calyx (ca) (D’-D’’’), with FasII and ChAT (magenta) labeling the neuropil. Scale bar: 50 µm. Abbreviations: ca (calyx), cb (central brain), ChAT (choline acetyltransferase), Dlg (Disc large), FasII (fasciclin II), mb (mushroom body), ml (medial lobe), pe (peduncle), sp (spur), vl (vertical lobe).

**Fig 3.**
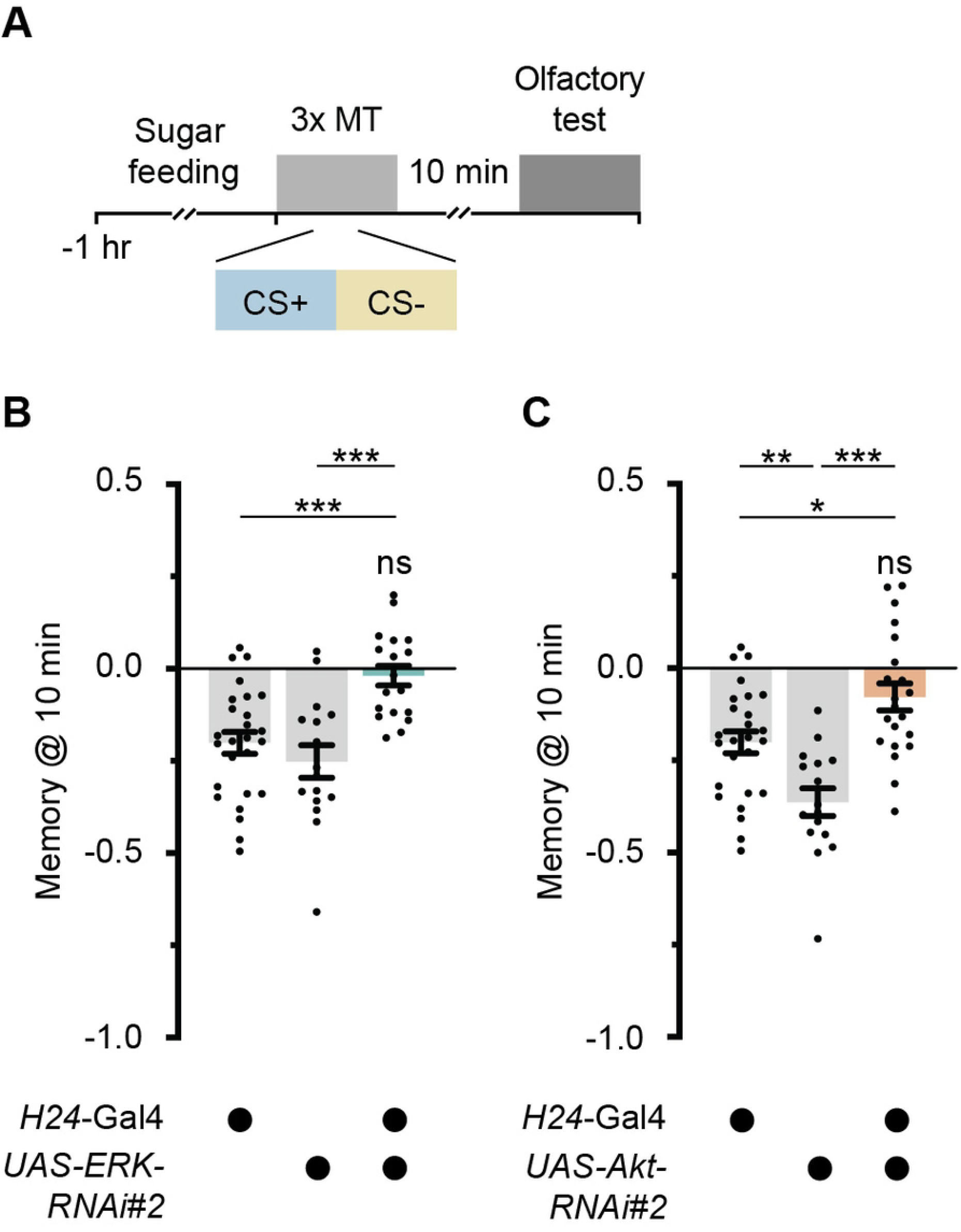
Impaired metabolically dependent memory from RNAi-mediated knockdown of ERK and Akt in larval mushroom bodies (MBs) (A) Larvae were fed sucrose for 60 minutes, and then subjected to three-cycle massed odor–high salt-conditioning using amyl acetate (AM) and benzaldehyde (BA), with one odor combined with high salt conditions. Larvae were memory-tested 10 minutes after training. (B) Larvae with RNAi-mediated downregulation of ERK in MB using *UAS-ERK-RNAi#2*, driven by the specific MB driver line *H24*-Gal4, exhibited total memory loss when tested for 10-minute memory. (C) Larvae with RNAi-mediated downregulation of Akt in the MB using *UAS-Akt-RNAi#2*, driven by *H24*-Gal4, showed total memory loss when tested for 10-minute memory. Data are shown as scatter dot plots, with each bar representing the mean ± SEM and each dot representing individual data points. Statistical significance was determined using one-way ANOVA and tested against the level of chance with a one-sample t-test or Wilcoxon signed-rank test. (***) p < 0.001, (**) p < 0.01, (*) p < 0.05, and ns, not significant. For further statistical details, see S3 Tables and S4 Table. Note, data for the *H24*-Gal4 control group shown in Figs. 3B and 3C were pooled from overlapping experiments.

## Results

### Localization of phosphorylated ERK and Akt in the larval central nervous system

In adult *Drosophila*, distinct regions of the MB are known to exhibit pERK, which is critical for memory formation [28,29,62]. Whereas the distribution of pERK in the *Drosophila* brain is well documented, less is known about the presence of phosphorylated Akt (pAkt) in the MB. Furthermore, the distributions of both kinases in the larval MB remain largely unexplored, despite the recognized multifunctionality of Akt and ERK in the larval brain [63]. To elucidate the roles of ERK and Akt in memory-related functions, we examined the distributions of pERK and pAkt, the active forms of these kinases, in the CNS of larval *Drosophila*.

Using anti-pERK and anti-pAkt antibodies with whole-mount brain preparations from 6-day-old larvae, and employing either the postsynaptic marker Discs large (Dlg) or a combination of anti-fasciclin II (FasII) and anti-choline acetyltransferase (ChAT) to identify neuropil areas within the larval CNS [64–66], we verified the presence of pERK throughout the larval CNS (Fig 1A-A’’), including the different MB areas (Fig 1B-D’’).

Similarly, pAkt was observed throughout the CNS (Fig 2A-A’’), with a notable presence within the MB (Fig 2B-D’’). The inactive (unphosphorylated) forms of ERK and Akt also showed a ubiquitous presence throughout the CNS and within the MB (S1A-C’’ Fig, S2A-C’’ Fig). The observed distributions of pERK and pAkt, along with their unphosphorylated counterparts, in the MB calyx and lobes suggests functional roles for these kinases in memory formation in *Drosophila* larvae. The specific presence of pAkt in the MB underscores its potential importance in these processes.

### Both kinases showed an impairment in metabolically dependent memory formation

Next, we analyzed the effectiveness of RNAi-mediated downregulation in various lines to investigate their roles in memory formation under elevated metabolic conditions. Using Western blot on adult fly heads with the pan-neuronal elav-Gal4 driver, we found that the UAS-ERK-RNAi#2 line (BDSC 34855) [54] significantly reduced pERK levels compared to UAS-ERK-RNAi#1 (VDRC 109108) [55] and the wild-type control (S3A,B Fig, S3 Table). The efficiency of Akt knockdown was assessed by an adult eclosion assay, which showed significant lethality with Akt downregulation pan-neuronal using the elav-Gal4 line. The UAS-Akt-RNAi#2 line (BDSC 33615) [54,55] exhibited more severe eclosion defects, with only 0.0% of adults emerging, compared to 68% in the UAS-Akt-RNAi#1 line (VDRC 103703) [55] and 96.7% in the wild-type control (S3C Fig, S3 Table). These results indicate distinct downregulation efficiencies and led us to select UAS-ERK-RNAi#2 and UAS-Akt-RNAi#2 for further studies. With this, we investigated their roles in memory formation under elevated metabolic conditions. A previous study indicated that high energy states induced by sugar feeding affect memory formation in a manner dependent on insulin receptor activity [42].

In line with these findings, we used a three-cycle massed odor–high-salt conditioning protocol in which larvae were fed sucrose for 60 minutes prior to training to elevate their energetic state (Fig 3A). The results revealed that larvae with disrupted ERK signaling in the MB, specifically targeted with the *H24*-Gal4 driver line (*H24*-Gal4>*UAS-ERK-RNAi#2*), showed no detectable memory (Fig 3B, S3 Table). By contrast, memory performance in control groups remained unaffected (Fig 3B, S3 Tables, S4 Table). This indicates that the memory formation facilitated by an energy surplus in control conditions is abolished when ERK signaling is inhibited. Similarly, downregulation of Akt in the MB led to no detectable memory in animals expressing *H24*-Gal4>*UAS-Akt-RNAi#2* (Fig 3C, S3 Table), whereas genetic controls exhibited measurable memory, albeit with significantly different scores (Fig 3C, S3 Tables, S4 Tables). The finding of significant impairment of 10-minute memory emphasizes the essential roles of ERK and Akt in the early stages of energy-induced memory consolidation.

### Morphological integrity of the mushroom body after ERK and Akt downregulation

Given the pivotal roles of ERK and Akt signaling during development [63], downregulation of ERK or Akt within the MB may lead to memory deficits due to a compromised MB structure. Therefore, we performed an immunohistochemical analysis to assess for any evident alterations in MB morphology.

Utilizing Dlg to identify the neuropil areas within the larval CNS, we focused on evaluating the morphological state of the following key MB areas: the lobes, peduncle, and calyx. Again employing the MB-specific *H24*-Gal4 driver line for targeted downregulation, we observed that neither *H24*-Gal4>*UAS-ERK-RNAi#2* (Fig 4A,A’) nor *H24*-Gal4>*UAS-Akt-RNAi#2* (Fig 4B,B’) resulted in discernible morphological alterations within the MB structures. Despite the targeted downregulation of Akt and ERK signaling pathways, key areas such as the lobes, peduncle, and calyx maintained their structural integrity, with no observable displacements or deformities (Fig 4A’,B’). This observation suggests that within the resolution limits of Dlg staining, the MB maintains its structural integrity despite reduced signaling activity from Akt and ERK. It’s important to note that more subtle changes or alterations not detectable by this method could potentially be uncovered with more sensitive techniques.

**Fig 4.**
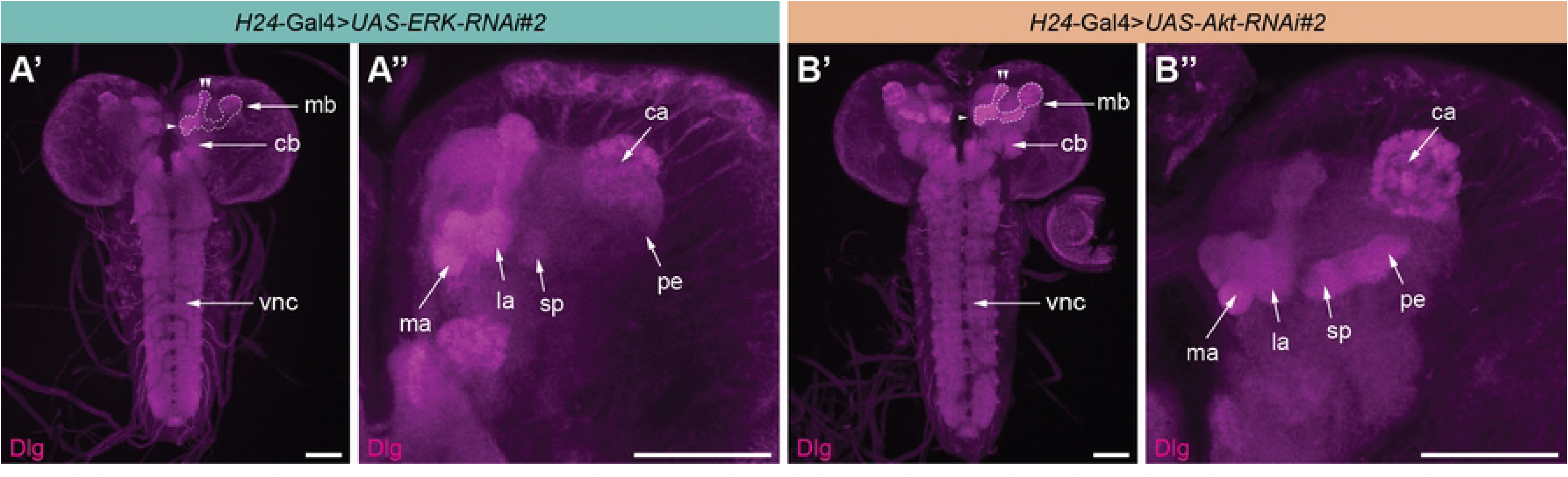
Normal mushroom body (MB) structure in larvae after RNAi-mediated knockdown of ERK and Akt. Maximum z-projections illustrate the central nervous system and mushroom body (MB) morphology in larvae following knockdown of ERK (A,A’) and Akt (B,B’), achieved using *UAS-ERK-RNAi#2* and *UAS-Akt-RNAi#2,* respectively, driven by the MB-specific *H24*-Gal4 driver. The calyx and lobes are highlighted in white. Anti-Dlg staining (magenta) provides a pan-neuronal anatomical backdrop. Scale bar: 50 µm. Abbreviations: ca (calyx), Dlg (Discs large), ma (medial appendix), la (lateral appendix), sp (spur), pe (peduncle). One arrowhead indicates the medial lobe (ml); two arrowheads indicate the vertical lobe (vl).

### Effects of downregulated ERK and Akt on naïve olfactory and gustatory perception

After assessing the structural integrity of the MBs in larvae with targeted downregulation of ERK and Akt, we investigated the functional outcomes of these manipulations. We specifically evaluated how RNAi-mediated reductions of ERK and Akt influenced sensory perception, which is essential to a comprehensive understanding of the roles of ERK and Akt in memory processes. We focused on responses relevant for odor-high salt conditioning namely the two odors amyl acetate (AM) and benzaldehyde (BA), as well as salt avoidance (Fig A’-A’’’).

The analysis revealed that larvae with downregulated ERK in the MB exhibited olfactory preferences and salt avoidance that were not statistically different from those of the genetic controls (Fig. 5B-D, S3 Tables, S4 Table). Similarly, larvae with downregulated Akt in the MB showed olfactory preferences and salt avoidance comparable to the genetic controls (Fig. 5E-G, S3 Tables, S4 Table). These results indicate that larvae expressing *H24*-Gal4>*UAS-ERK-RNAi#2* and *H24*-Gal4>*UAS-Akt-RNAi#2* retain sensory perception that are comparable to those of the genetic control groups. Despite the targeted downregulation of ERK and Akt within the MBs, these critical aspects of sensory function appear to remain intact and suggest that the specific roles of ERK and Akt do not extend to impairing fundamental sensory processes.

**Fig 5.**
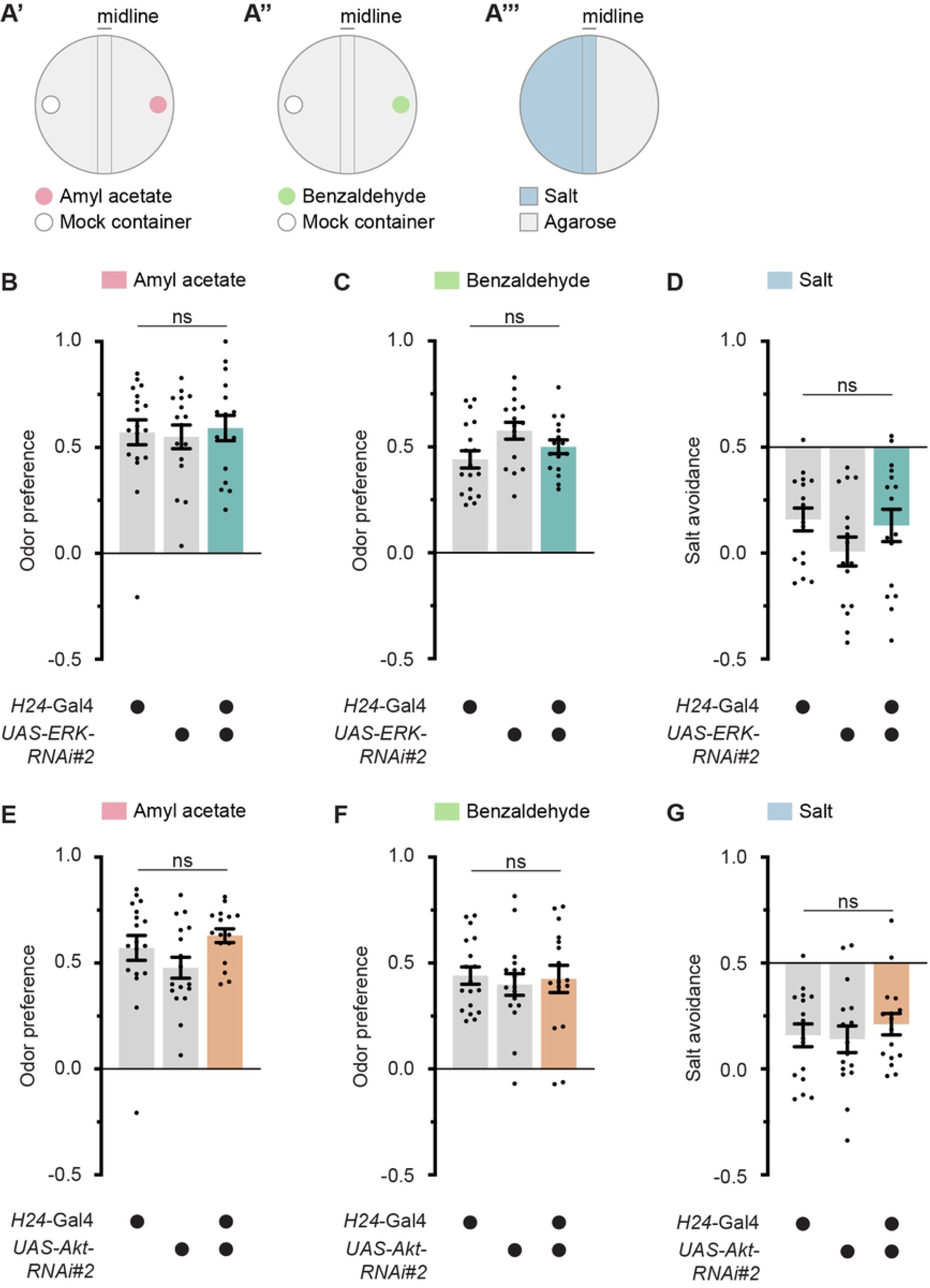
Naïve olfactory preferences and salt avoidance in larvae with RNAi-mediated ERK and Akt knockdown in the mushroom body (MB) (A’-A’’’) Schematic drawing of naïve odor preference and high-salt avoidance assays. For odor preference, larvae were placed along the midline of an agarose-filled Petri dish. Amyl acetate (AM, 1:250) (A’) or benzaldehyde (BA, undiluted) (A’’) was placed on one side, with a mock container on the other. For high-salt avoidance (A’’’), one side contained agarose and the other agarose with 1.5 M sodium chloride (salt). Larval distribution was evaluated after 5 minutes. (B-D) Knockdown of ERK in the mushroom body (MB) using *UAS-ERK-RNAi#2* driven by the MB-specific *H24*-Gal4 driver did not affect naïve responses to AM (A), BA (B), or salt (C) compared to both control groups (*H24*-Gal4/+ and *UAS-ERK-RNAi#2*/+). (E-G) Similarly, knockdown of Akt in the MB using *UAS-Akt-RNAi#2* driven by *H24*-Gal4 evoked no differences in naive responses to AM (A), BA (B), or salt (C) compared to both control groups (*H24*-Gal4/+ and *UAS-Akt-RNAi#2*/+). Data are shown as scatter dot plots, with each bar representing the mean ± SEM and each dot representing individual data points. Statistical significance was determined using one-way ANOVA or Kruskal-Wallis test. (***) p < 0.001, (**) p < 0.01, (*) p < 0.05, and ns, not significant. For further statistical details, see S3 Tables and S4 Table. Note, data for the *H24*-Gal4 control group shown in Figs. 5B and 5E, 5C and 5F, and 5D and 5G were pooled from overlapping experiments.

### No detectable effects of downregulated ERK and Akt on associative olfactory learning

To further understand the role of ERK and Akt in memory processes, we tested whether downregulation of these kinases affects olfactory learning and memory, independent of the energy state.

Using a single-cycle odor–high-salt conditioning protocol as described [5] (Fig 6A), we assessed immediate memory to determine if basic learning processes were impaired. Notably, larvae expressing *H24*-Gal4>*UAS-ERK-RNAi#2* demonstrated immediate memory scores that were not statistically different from those of either genetic control (*H24*-Gal4/+ and UAS-ERK-RNAi#2/+) (Fig 6B, S3 Table, S4 Table). Similarly, larvae expressing *H24*-Gal4>*UAS-Akt-RNAi#2* also showed immediate memory scores that matched the performance of both genetic controls (*H24*-Gal4/+ and *UAS-Akt-RNAi#2*/+) (Fig 6C, S3 Table, S4 Table). In summary, despite the targeted downregulation of ERK and Akt within the MB, critical aspects of basic olfactory learning and memory processes appear to remain intact and suggest specific roles for ERK and Akt in metabolically dependent memory formation processes.

**Fig 6.**
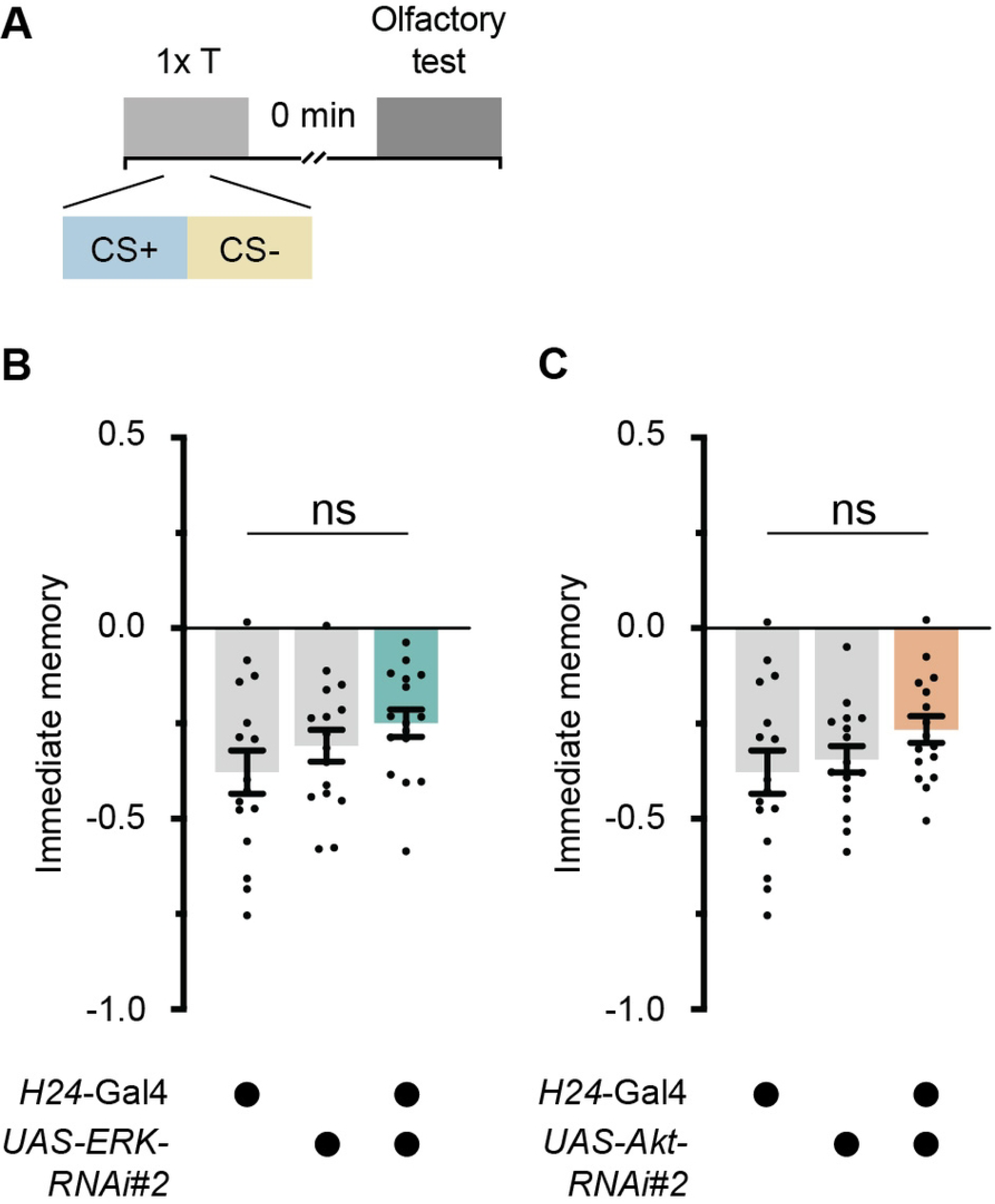
Impact of RNAi-mediated knockdown of ERK and Akt in larval mushroom bodies (MBs) on immediate olfactory memory. (A) Larvae underwent single-cycle odor–high-salt conditioning using amyl acetate (AM) and benzaldehyde (BA), with one odor combined with the high-salt condition. Memory was tested directly after training (immediate memory). (B) Larvae with a knockdown of ERK (*H24*-Gal4>*UAS-ERK-RNAi#2*) showed immediate memory that was indistinguishable from the genetic control groups (*H24*-Gal4/+ and *UAS-ERK-RNAi#2*/+). (C) Larvae with a knockdown of Akt (*H24*-Gal4>*UAS-Akt-RNAi#2*) showed immediate memory that was indistinguishable from genetic control groups (*H24*-Gal4/+ and *UAS-Akt-RNAi#2*/+). Data are shown as scatter dot plots, with each bar representing the mean ± SEM and each dot representing individual data points. Statistical significance was determined using one-way ANOVA or Kruskal-Wallis test. (***) p < 0.001, (**) p < 0.01, (*) p < 0.05, and ns, not significant. For further statistical details, see S3 Table and S4 Table. Note, data for the *H24*-Gal4 control group shown in Figs. 6B and 6C were pooled from overlapping experiments.

## Discussion

Previous research in *Drosophila* has established that certain biological and environmental conditions must be met for LTM to form. Typically, memory formation involves spaced training that allows for critical transcriptional activities essential for memory consolidation, such as the interplay between CREB and the immediate early gene c-Fos [4–6,28,62,67–69]. However, emerging studies suggest that these traditional requirements can be modified by various factors—including metabolic state [27–29,42]. For instance, insulin receptor activity, which is influenced by elevated energy levels, has been shown to modulate memory processes and allow for the formation of LTM without the need for temporal spacing [42].

In this study, we expanded on these findings by analyzing the roles of downstream targets of the insulin receptor—specifically ERK and Akt kinases—in metabolically-dependent memory formation. We employed a protocol in which animals were fed sugar for 60 minutes before undergoing a three-cycle massed odor–high-salt conditioning regime, which facilitates sugar-dependent LTM formation [42]. Our results showed that reducing ERK and Akt levels in the MB significantly impaired memory under conditions of increased energy states, observable as soon as 10 minutes after training (Fig. 4). This suggests that ERK and Akt are directly involved in memory formation, rather than regulating a gating process between different types of memory, as previously shown with the insulin receptor acting upstream of both kinases [42].

We hypothesize that the insulin receptor serves as a gating mechanism in high-energy states, and that it fulfills two critical functions. Initially, it inhibits the formation of another consolidated memory type that is not dependent on protein synthesis through the suppression of Rho kinase (ROCK), mediated by Ras-Raf signaling, which is a downstream target of the insulin receptor [29,52]. Concurrently, it activates ERK and Akt, both of which are known to be crucial regulators of memory formation in various species [28,62,70–79], which then mediate the formation of metabolically dependent memory (Fig. 4). Notably, ERK phosphorylation during the rest intervals in spaced training protocols is well documented, suggesting a pivotal role, not only in the consolidation process but also in the early stages of LTM formation [28,52]. We observed learning and memory deficits that further support the critical early activation of ERK, essential for initiating rapid responses to sugar-induced LTM formation. This early activation could also be paralleled by Akt, although the specific timing of Akt activation remains less defined. The early and rapid activation of both ERK and Akt is crucial to consolidate metabolically dependent LTM. However, further studies are needed to disentangle which specific aspects of memory formation are affected by this impairment, and to what extent they are necessary for energy-induced LTM compared to their roles in training-induced LTM.

An alternative explanation could be that the constant downregulation of ERK and Akt compromised the neuronal architecture of the MB, given their known functions during embryonic and larval development [63]. Although developmental defects were ruled out by confirming the morphological integrity of the MB (Fig. 5), the potential impact of downregulation on neuronal circuitry requires further investigation. Our tests confirmed that naïve olfactory and gustatory perceptions were unchanged following the downregulation of ERK and Akt in the MB (Fig 6), suggesting that the observed memory impairments were not due to sensory deficits. Furthermore, the impairment was specific to metabolically dependent memory, as learning itself was not affected (Fig 6).

In conclusion, our study expands upon the traditional view that memory formation is primarily governed by neuronal circuitries, such as serotonergic and dopaminergic input to the MB [26,27], by emphasizing the critical role of molecular signals within the MB. Our findings align with previous work showing that molecular pathways, particularly those involving insulin receptor signaling through ERK and Akt, are critical for memory processes [28,29,42]. Ultimately, our research suggests a connection between metabolically dependent memory formation and the complex interplay of neuronal and molecular signaling. These findings not only enhance our understanding of memory processes in *Drosophila,* but also suggest a potentially broader biological significance of ERK and Akt signalling in memory processes across species.

## Acknowledgments

We are grateful to André Fiala for his support. We thank to Bele Metzner for contributing to experiments. We also thank the Bloomington Drosophila Stock Center (BDSC) and the Vienna Drosophila Resource Center (VDRC) for providing fly lines. This research was supported by the Deutsche Forschungsgemeinschaft (DFG, German Research Foundation) under grant number 459524845 to AW.

## Supporting information

**S1 Fig.**
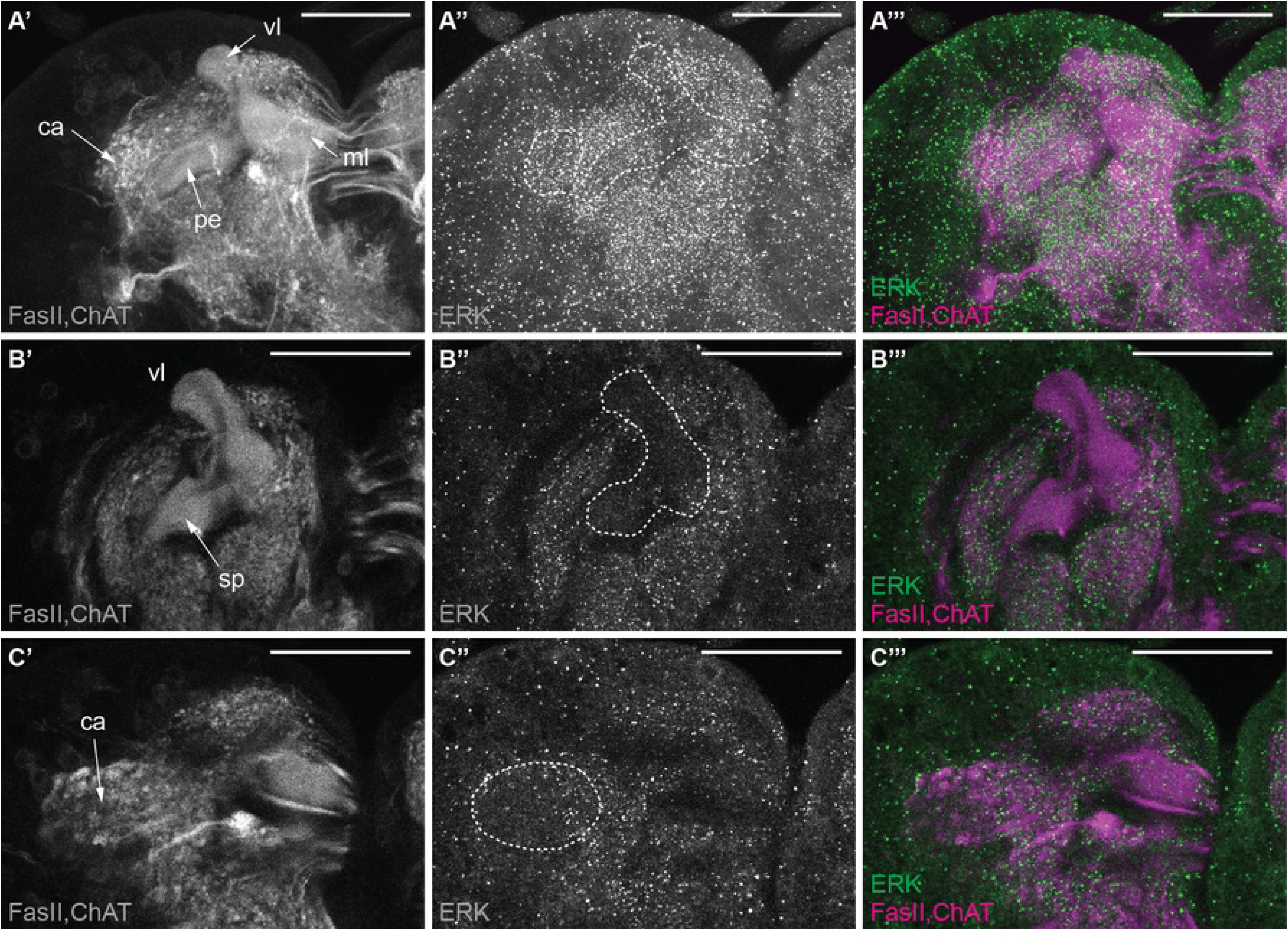
Localization of unphosphorylated ERK in the mushroom body (MB) (A’-A’’’) Maximum z-projection of unphosphorylated ERK localization (green) in the MB, with FasII and ChAT (magenta) highlighting the neuropil. (B’-C’’’) Single-slice views highlight the distribution of pAkt in MB subregions, including visible portions of the lobes (vl, s) (B’-B’’’) and calyx (ca) (C’-C’’’), with FasII and ChAT (magenta) labeling the neuropil. Scale bar: 50 µm. Abbreviations: ca (calyx), ml (medial lobe), pe (peduncle), sp (spur), vl (vertical lobe).

**S2 Fig.**
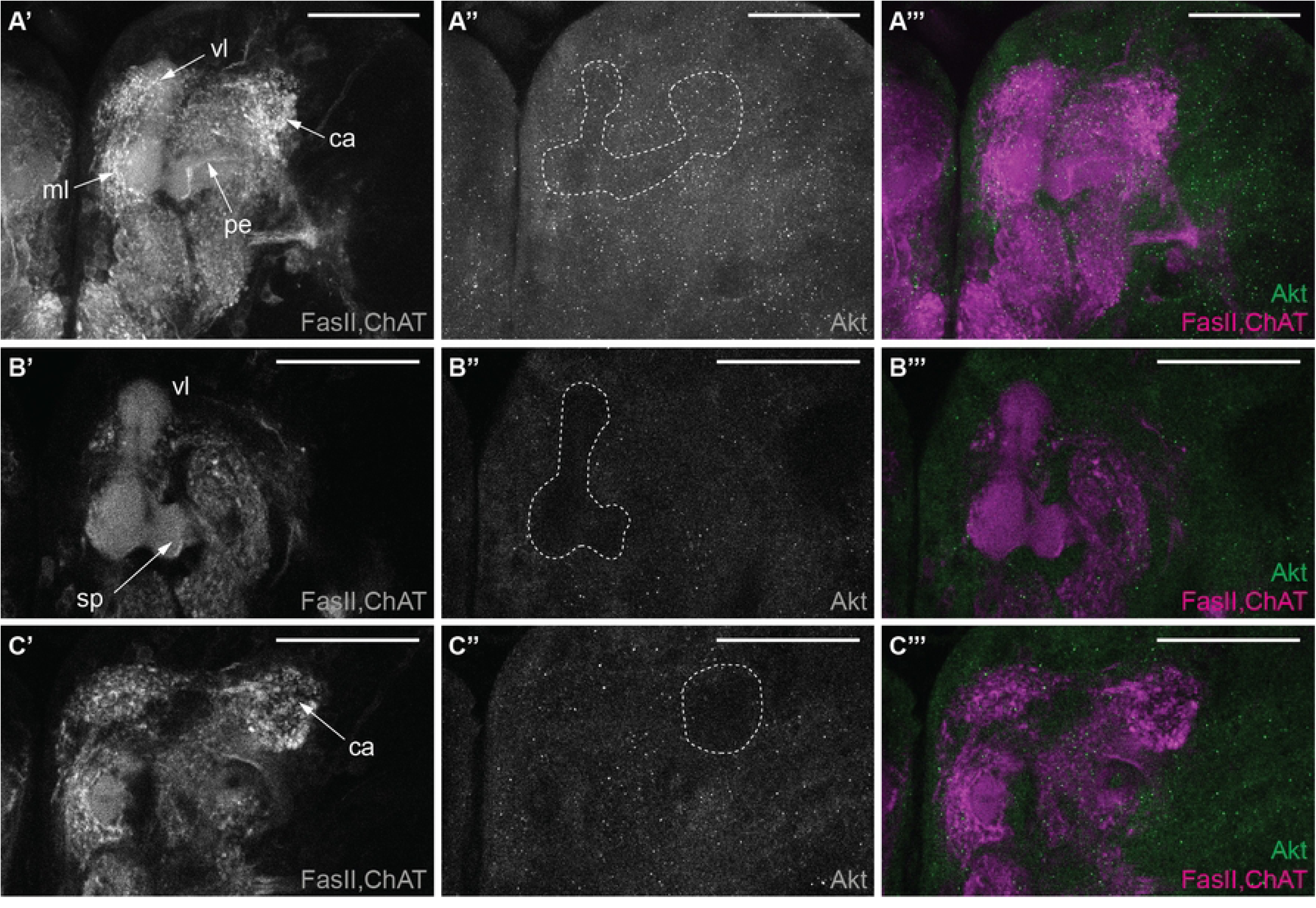
Localization of unphosphorylated Akt in the mushroom body (MB) (A’-A’’’) Maximum z-projection unphosphorylated Akt visualization (green) in the MB lobes, with FasII and ChAT (magenta) highlighting the neuropil. (B’-C’’’) Single-slice views highlight the distribution of pAkt in MB subregions, including visible portions of the lobes (vl, s) (B’-B’’’) and calyx (ca) (C’-C’’’), with FasII and ChAT (magenta) labeling the neuropil. Scale bar: 50 µm. Abbreviations: ca (calyx), ml (medial lobe), pe (peduncle), sp (spur), vl (vertical lobe).

**S3 Fig.**
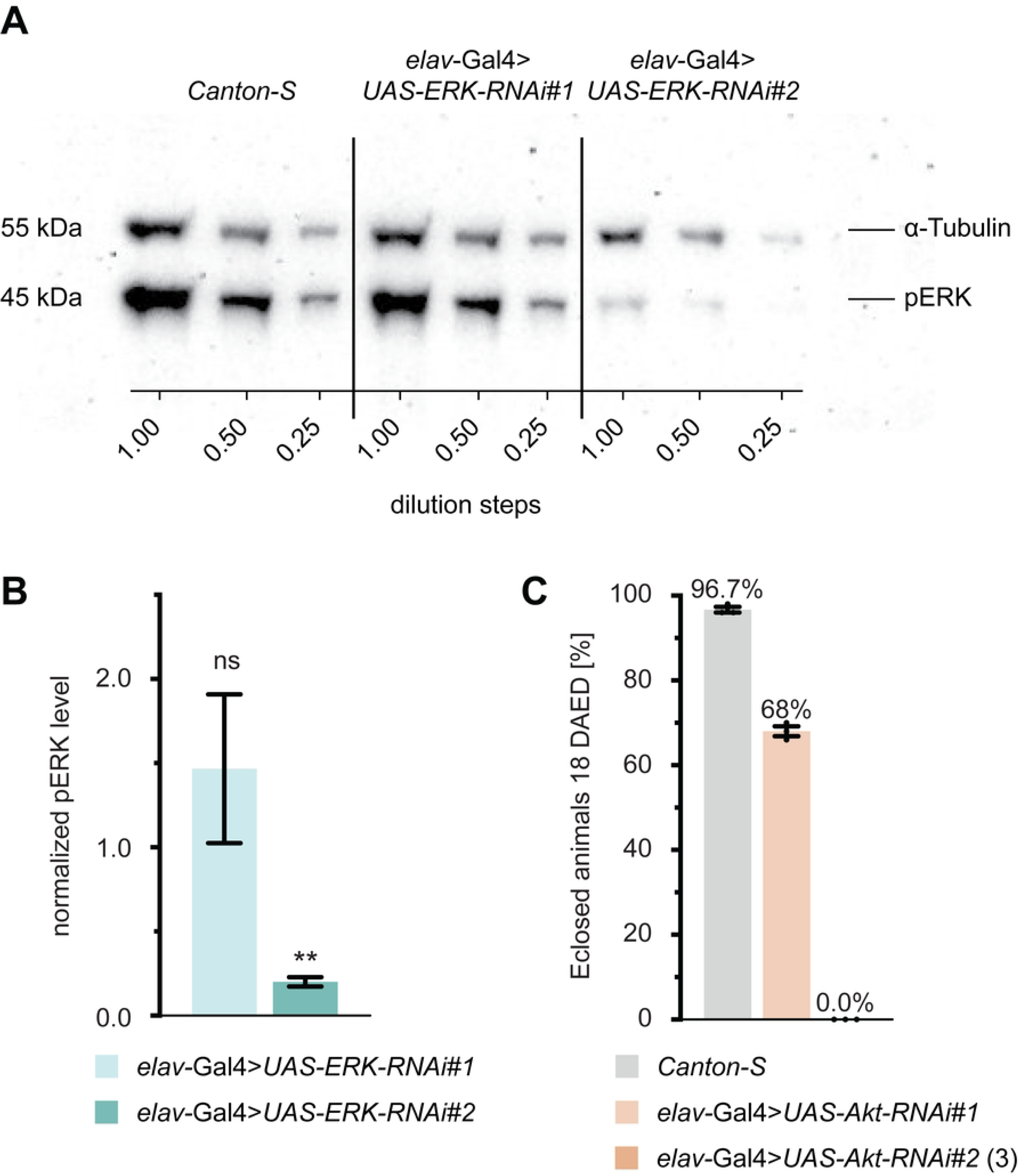
Assessment of ERK and Akt knockdown efficacy in *Drosophila*. (A) Evaluation of ERK phosphorylation levels following knockdown in the adult Drosophila central nervous system (CNS). Western blot of pERK levels in head lysates from adult flies, with ERK knockdown achieved pan-neuronally using two RNAi lines (*UAS-ERK-RNAi#1* and *UAS-ERK-RNAi#2*) driven by *elav-*Gal4. α-Tubulin (α-Tub) served as an internal loading control for protein quantification. (B) Normalized pERK levels demonstrate a significant reduction in *elav*-Gal4>*UAS-ERK-RNAi#2* fly heads (0.20 ± 0.04) but not in *elav*-Gal4>*UAS-ERK-RNAi#1* (1.47 ± 0.59). Bar plots depict the mean normalized pERK relative to wild-type control values from three Western blots, with SEM error bars. A one-sample t-test was performed to evaluate if the normalized pERK levels significantly deviated from a theoretical mean of μ=1, corresponding to normalized wild-type control levels. (***) p < 0.001, (**) p < 0.01, (*) p < 0.05, and ns, not significant. (C) Eclosion rates of *Drosophila* after pan-neuronal downregulation of Akt in the CNS. *Canton-S* (wild-type control) flies have an eclosion rate of 96.7%, *elav*-Gal4>*UAS-Akt-RNAi#1* flies have an eclosion rate of 68%, and *elav*-Gal4>*UAS-Akt-RNAi#2* flies have an eclosion rate of 0.0%. Bar plots depict the mean eclosion rate in percentage, with SEM error bars, taken from 18 days after egg deposit (DAED). For further statistical details, see Tables 3.

**S1 Table. *Drosophila* strains used in this study.**

**S2 Table. Key resource tables.**

**S3 Table. Sample size, mean±s.e.m. and statistical details of test against chance level.**

**S4 Table. Statistical details of ordinary one-way ANOVA and non-parametric Kruskal-Wallis test.**

